# Sodium-enriched nectar increases seed set in low diversity prairies

**DOI:** 10.64898/2026.11.25.769002

**Authors:** Ashley L. Darst, Megan E. Lamm, Nick M. Haddad

## Abstract

Plant-pollinator interactions depend on floral traits and background plant community properties, yet the role of nectar micronutrients in these interactions is poorly understood. Sodium is a micronutrient of particular interest because although plants need little to no sodium, animal pollinators require it. It is unknown whether sodium-enriched nectar increases plant fitness in animal-pollinated plants and how this may be affected by the broader plant community. To test this, we grew black-eyed Susan (*Rudbeckia hirta*) plants in sodium-enriched or control soil. We placed pairs of sodium-addition and control plants in prairies seeded with high or low diversity and measured seed set and germination rate. In a supplementary experiment, we explored differences in floral visitation. We found that black-eyed Susan plants grown in sodium-enriched soil had sodium-enriched nectar. Seed set increased in sodium-addition plants in prairies seeded with low diversity but not in prairies seeded with high diversity. Germination rate was not affected by sodium addition. Altogether, our study highlights the importance of considering both micronutrients and context dependency in understanding plant reproduction and complex species interactions.

## Background

Plant-pollinator interactions depend on many factors, including both the properties of the interacting species and the properties of the communities they inhabit [1,2]. Floral nectar is one important factor as a reward for pollinators, with traits such as nectar volume, sugar concentration, and composition influencing interactions [3,4]. However, the role of nectar micronutrients in shaping plant-pollinator interactions is poorly understood [5–7]. Sodium is a micronutrient of particular interest because plants and pollinators differ in their sodium requirements. While plants need little to no sodium, animals require it for essential bodily functions such as maintaining homeostasis and generating nerve impulses [6,8–10]. In sodium-limited continent interiors [11,12], some pollinators display sodium-seeking behaviors to meet their dietary requirements from sodium-rich mineral and animal sources [13–16]. However, many pollinators have not been documented “puddling” and may instead acquire sodium through plant material [17,18]. One possible source of sodium in plants is nectar [19–21]. The ‘salty nectar hypothesis’ posits that if plants can increase the sodium concentration in their nectar and attract more or higher quality pollinators, they might outperform those that cannot [6]. However, plants are not isolated in nature and pollinator preference for sodium-enriched nectar may depend on the properties of the background plant community [22,23]. It is unknown whether most plants can transport sodium into their nectar and whether this increases plant fitness in a variety of plant communities.

For sodium to increase plant fitness via sodium-enriched nectar, sodium should be transported into nectar and be preferred by pollinators. Recent studies found twice as many floral visitors to plants with sodium-enriched nectar [24–26], but the consequences of this increased visitation on plant fitness are unknown. Pollinator quantity and quality are both important to plant fitness [27]. For example, more floral visitors may not increase plant fitness if they are nectar robbers, nectar thieves, or pollen thieves [28,29], or if the floral visitors are lower quality pollinators [27]. Additionally, unless plants can actively transport sodium from foliar tissue into nectar, increased foliar sodium from elevated soil sodium may reduce fitness through increased herbivory [6,30].

The background plant community is also important to consider for pollinator behavior and plant fitness. In particular, plant community diversity may add context dependency to the ‘salty nectar hypothesis’. While greater floral diversity attracts a greater abundance and diversity of pollinators [31,32], pollinators may not be able to detect or remember the locations of favored rewards in a plethora of colors, shapes, and floral volatiles [33–36]. Therefore, even if pollinators prefer plants with sodium-enriched nectar, pollination may depend on the broader floral community diversity.

In this study, we compared seed set and germination rate as components of fitness for black-eyed Susan (*Rudbeckia hirta*) plants grown in sodium-enriched and control soil in prairies seeded with high and low diversity. In a supplementary experiment to investigate pollinator preference for sodium-enriched nectar, we additionally measured aspects of floral visitation to sodium-enriched and control plants. Black-eyed Susan plants are obligate outcrossers that are tolerant of high soil sodium concentrations [37,38], and they have been previously used in studies investigating pollination rate [39,40]. We measured both seed set and germination rate to investigate how sodium could affect multiple components of plant fitness [41]. Following the ‘salty nectar hypothesis’ [6], we predicted that sodium-enriched nectar would increase floral visitation and seed set in the animal-pollinated black-eyed Susan. We further predicted that the effect of sodium on seed set would be enhanced in plant communities with lower diversity.

## Methods

### Greenhouse methods

We propagated black-eyed Susan (*Rudbeckia hirta* L.) plants in control or sodium-enriched soil. We sourced black-eyed Susan seeds from Iowa (Prairie Moon Nursey). We planted the seeds at W.K. Kellogg Biological Station (Hickory Corners, Michigan, USA) in a 21.1°C greenhouse with a 16/8 hour light/dark cycle at the end of February 2024. We fertilized seedlings with a low dose of NPK fertilizer and transplanted them into one-gallon pots. We added a total of 7.5 g of NaCl to the soil of a randomly chosen half of the plants over four weeks. We chose this amount of sodium to mimic sodium concentrations found in roadsides where sodium deicers are used and that are also tolerable to black-eyed Susan plants [37,42,43]. For the first three applications (one per week), we dissolved 1.5 g NaCl in 200 mL water and applied the brine directly to the soil. For the last application before deployment to the field, we applied 3 g NaCl dissolved in 200 mL water. To the other half of the plants, we applied 200 mL water to the soil per week concurrently with sodium brine applications. We placed trays under pots to prevent sodium running through. We watered plants as needed outside of applications, with all plants receiving approximately the same amount of water during each application.

### Sodium analysis

To understand sodium movement in black-eyed Susan plants, we quantified sodium in the nectar, leaves, and soil in greenhouse plants not deployed into the field that received sodium-addition and control treatments (n = 6 plants per treatment). We quantified nectar sodium using two methods: the centrifuge method and the rinse method. For the centrifuge method, we harvested five flower heads per plant with open florets and trimmed them for easier access to the nectar [44]. We centrifuged each flower head through a fine mesh into an Eppendorf Tube at 3500 rpm (∼1500 g) for ten minutes, which is unlikely to cause cell rupturing [45]. After all five flower heads’ nectar was centrifuged into the same tube, we used 15 µL of ultrapure type 1 water to rinse the nectar off the side of the tube and collected 10 µL of nectar/water solution. There was likely some pollen contamination using this centrifuge method, and we consider this possibility in our interpretation of the results. For the rinse method, we harvested an additional two flower heads per plant and stirred each flower head individually in 2 mL of ultrapure type 1 water for two minutes [46]. This rinse method included both nectar and pollen. Unfortunately, our methods of nectar extraction prevent us from making meaningful comparisons between the concentration of sodium in the foliar tissue and nectar. Therefore, we cannot infer whether transport of sodium into the nectar was passive or active. We collected leaves and soil from each plant on the same day. Approximately ten leaves were collected per plant, rinsed with DI water, and dried in a drying oven at 57°C for at least 48 hours. We used a hand trowel to sample soil for the entire depth of the one-gallon pot.

Sodium quantification was performed at the Quantitative Bio Element Analysis and Mapping center at Michigan State University using ICP-QQQ-MS. For centrifuged nectar, samples were digested in 200 µL HNO_3_ and heated at 65°C for 2 hours. Samples received 300 µL more HNO_3_ and were diluted up to 10 mL with water. For nectar and pollen from the rinse method, samples were vortexed and 500 µL were digested in 500 µL HNO_3_ and heated at 65°C for 2 hours. Samples were diluted up to 10 mL with water. For leaf analyses, between 25–50 mg of leaf powder were weighed out per sample and digested in 2.5 mL HNO_3_ using microwave assisted acid digestion protocol 3051 [47]. Samples were diluted up to 50 mL with water. For soil analyses, between 20–30 mg soil were weighed out per sample and digested in HNO_3_. Samples were diluted up to 10 mL with water.

### Field site

To investigate the effects of plant community diversity on the floral visitation, seed set, and germination rate of sodium-enriched and control black-eyed Susan plants, we used experimentally restored prairies in the Conservation Lands Experiment (CLE) at the W.K. Kellogg Biological Station (42.4°, −85.4°) [48]. This region is naturally sodium-poor [6,49]. The CLE was formerly agricultural land and was seeded with prairie species in 2015. The CLE has 12 prairie plantings ranging from 0.2–3.5 hectares with half of each seeded with a high diversity seed mix (70 seeded species, including 50 forbs) and the other half seeded with a low diversity seed mix (12 seeded species, including 7 forbs). Both seed mixes included black-eyed Susan (*Rudbeckia hirta*), but they are not common in the established prairie communities. Previous studies have found greater species richness in the high diversity treatment at the whole-field scale [48]. There already existed two permanent herbivore exclosures within each diversity treatment replicate (total = 48 exclosures) [50]. One of the two herbivore exclosures in each replicate was located near the center of the planting while the other was located near the edge. We deployed plants within these herbivore exclosures to prevent herbivory from large mammals present at our site that are known to consume experimentally watered plants. Herbivory from insects was still possible.

### Germination and seed set

To test the effects of sodium enrichment and plant community diversity on black-eyed Susan seed set and germination rate, we deployed 96 plants in control/sodium-addition pairs into the herbivore exclosures in the CLE. In the greenhouse just prior to deployment to the field, one flower per plant not yet producing pollen was enclosed in a pollinator exclusion bag [40]. We placed one control and one sodium-addition plant within each exclosure and removed the pollinator exclusion bags. Plants remained in the field for 17 full days (not including the days deploying and removing plants). The first plants were deployed May 31, 2024 and the last plants were removed June 20, 2024. After 17 days, pollinator exclusion bags were replaced on the flower heads open to pollination. All plants were moved to a common location under a shade cloth until the seed heads matured. Seed heads were harvested once fully dry, between July 25, 2024 and August 13, 2024. Seed heads were stored in a refrigerator.

In January 2025, we randomly chose thirty achenes (“fruits”) from each seed head for a germination trial [40]. We randomly selected achenes regardless of any apparent seed development using a numbered grid with a random number generator to select the grid cell. We placed achenes in a petri dish on a damp paper towel and germinated them in 21.1°C growth chambers with a 16/8 hour light/dark cycle. We checked achenes daily for germination and kept them damp. After 40 days, we stopped the germination trial. We calculated germination rate as the number of germinated seeds out of the total number of developed seeds (as determined by either dissection or germination). The total number of developed seeds was often less than thirty.

To determine seed set, we dissected the remaining ungerminated achenes. Achenes containing embryo were considered seeds while empty achenes were not. We calculated seed set as the number of seeds (as determined by either dissection or germination) out of the total number of achenes (i.e., thirty).

### Floral visitation surveys

In a supplementary experiment to further understand how sodium-enriched nectar affected plant fitness, we conducted floral visitation surveys the following summer to explore visitor preference (see Supplementary Materials for detailed methods).

### Statistical analyses

Before we tested our core hypotheses, we conducted analyses of sodium concentration in the soil, leaves, and nectar in control and sodium-addition black-eyed Susan plants. We performed natural logarithm transformations of soil, leaf, and nectar concentrations prior to analyses because concentrations are often log-normal and transformations improved our model diagnostics [51]. For soil and leaf sodium concentrations, we used linear models with the natural logarithm of sodium concentration in µg/g as a function of sodium treatment (R package ‘stats’ v.4.6.1). Eight out of thirty-six nectar sodium content values were below the limit of detection (LOD ∼ 100 ng). We chose to include the values below the LOD in our models, instead of using imputation or censoring, because including these values often improves estimates [52]. Using a linear mixed effects model in R package ‘lme4’ v.2.0.1 [53], we took the natural logarithm of nectar sodium content in ng as a function of sodium treatment and sampling method. We included plant as a random intercept to account for repeated sampling. A censored regression model using the above formula without random effects yielded similar results [54]. Censored regression models including plant identity as a random effect did not converge.

To further understand how sodium moved within plants, we used structural equation modeling (SEM) using the function ‘sem’ in R package ‘lavaan’ v.0.7.2 [55]. We fit three models to be included in the SEM: 1) the natural logarithm of soil sodium (µg/g) as a function of sodium treatment, 2) the natural logarithm of leaf sodium (µg/g) as a function of the natural logarithm of soil sodium (µg/g), and 3) the natural logarithm of nectar sodium (ng/sample) as a function of the natural logarithm of leaf sodium (µg/g). Only centrifuged nectar samples were used for the SEM. We plotted the resulting path diagram using the function ‘semPaths’ in R package ‘semPlot’ v.1.2.0 [56].

To test the hypothesis that sodium increases seed set, we fit a generalized linear mixed effect model using a binomial distribution in R package ‘lme4’ v.2.0.1 [53]. We fit seed set as a function of sodium treatment with pair nested in CLE site as a random intercept. The best fitting and most parsimonious model (dAICc > 2, Table S2) supported our decision to include an interaction with seeded diversity (high/low). Post-hoc comparisons were performed on significant interactions in R package ‘emmeans’ v.2.0.3 [57], and p-values were adjusted using the Tukey method.

To test whether sodium negatively affected germination as a second component of fitness [58,59], we fit a generalized linear mixed effect model using a binomial distribution in R package ‘lme4’ v.2.0.1 [53]. We fit germination rate as a function of sodium treatment with the random intercepts pair nested in CLE site and growth chamber shelf nested in growth chamber. The best fitting and most parsimonious model (dAICc > 2, Table S2) with sodium treatment included seeded diversity (high/low) and exclosure location (edge/middle of field) as fixed effects.

All analyses were performed in R version 4.6.1 [60]. Linear models were compared using the AICctab function in R package ‘bbmle’ v.1.0.25.1 [61]. Linear model assumptions were assessed using diagnostic plots with R package ‘performance’ v.0.17.1 [62]. Test statistics and p-values were calculated using type II ANOVA for additive models and type III ANOVA for models with interaction terms in R package ‘car’ v.3.1.5 [63]. Regression coefficients from models with log-transformed response variables were back-transformed to percent changes using (exp(*β̂*) − 1) × 100.

## Results

After sodium addition, black-eyed Susan plants had elevated sodium in the soil, leaves, and nectar (Figure 1). On average, there was 1499% more µg sodium per gram soil for sodium-addition plants than control plants (*β̂* ± SE = 2.77 ± 0.13, F_1,10_ = 422.44, p < 0.0001). On average, there was 4212% more µg sodium per gram leaf for sodium-addition plants than control plants (*β̂* ± SE = 3.76 ± 0.72, F_1,10_ = 27.46, p = 0.0004). On average, there was 282% more ng sodium in nectar samples taken from sodium-addition plants than control plants (*β̂* ± SE = 1.57 ± 0.77, χ^2^ = 4.14, df = 1, p = 0.04), regardless of the sampling method used (rinse: *β̂* ± SE = −1.19 ± 0.74, χ^2^ = 2.60, df = 1, p = 0.11). The corresponding SEM fit the data well (CFI = 0.982, TLI = 0.963, SRMR = 0.028). Soil sodium concentration increased in sodium-enriched treatments (estimate ± SE = 2.77 ± 0.12, p < 0.001). Leaf sodium concentration increased with greater soil sodium concentration (estimate ± SE = 1.29 ± 0.26, p < 0.001). Nectar sodium content increased with greater leaf sodium concentration (estimate ± SE = 0.571 ± 0.25, p = 0.024).

**Figure 1.**
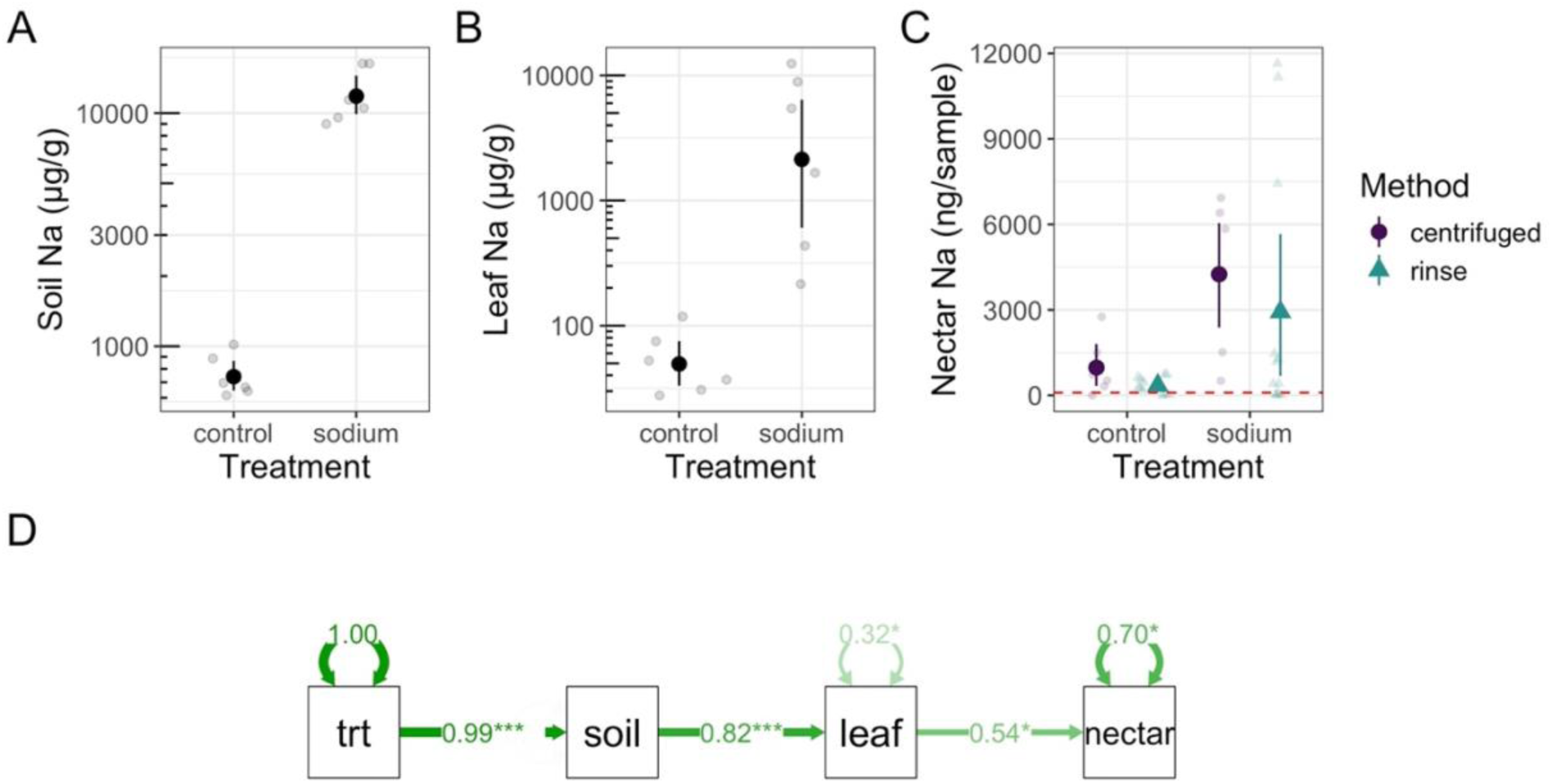
Sodium content in the A) soil, B) leaves, and C) nectar of sodium-addition and control black-eyed Susan plants. Solid points represent the mean. Error bars are 95% bootstrapped confidence intervals. More transparent points are raw, jittered data. The red dashed line represents the approximate limit of detection (100 ng/sample). D) The corresponding path diagram for sodium transport in black-eyed Susan plants. Values in weighted straight arrows are standardized parameter estimates and values in weighted circular arrows are residuals. Path diagram nectar values are from centrifuged samples only. *p-value < 0.05, **p-value < 0.01, ***p-value < 0.001.

Seed set increased in sodium-addition plants in prairies seeded with low diversity but not in prairies seeded with high diversity (Figure 2A, Table S7). There was a significant interaction between sodium treatment and seeded diversity (χ^2^ = 10.51, df = 1, p = 0.001). In low diversity prairies, sodium-addition plants had 40% greater odds of seed set than control plants (odds ratio ± SE = 1.397 ± 0.176, p = 0.040). For control plants, plants in high diversity prairies had 118% greater odds of seed set than those in low diversity prairies (odds ratio ± SE = 2.184 ± 0.63, p = 0.034). All other pairwise comparisons were non-significant (p > 0.19, Table S8).

**Figure 2.**
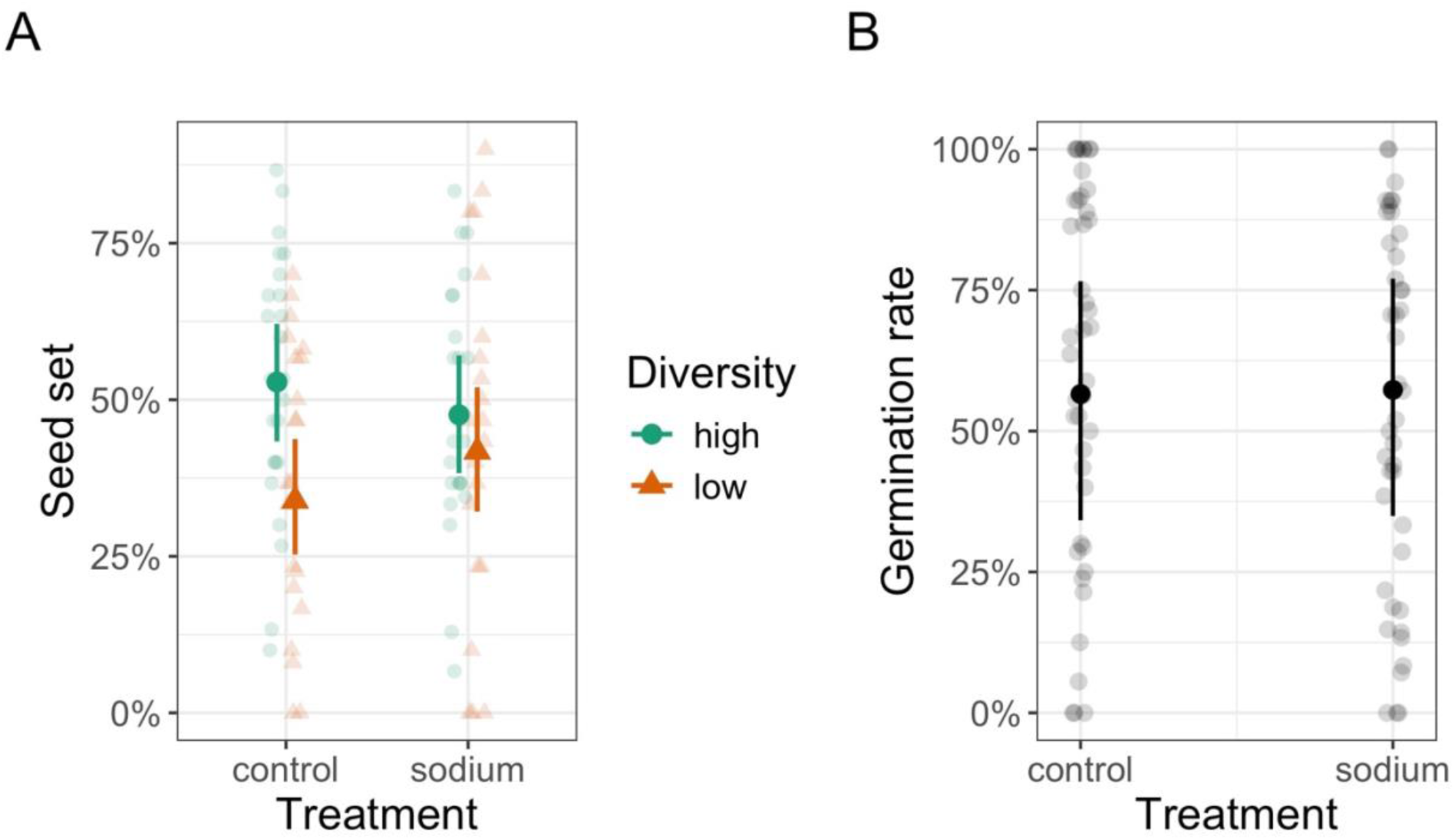
A) Seed set and B) germination rates for sodium-addition and control black-eyed Susan plants. The interaction of sodium treatment with high and low seeded diversity is shown for seed set. Solid points represent the estimated marginal means. Error bars are 95% confidence intervals. Translucent points are raw, jittered data.

Sodium-addition did not affect germination rate (χ^2^ = 0.03, df = 1, p = 0.854, Figure 2B, Table S7). The odds of germinating increased by 116% in high diversity prairies (χ^2^ = 3.95, df = 1, p = 0.047) and by 145% in exclosures near the middle of prairies (χ^2^ = 5.30, df = 1, p = 0.021, Figure S1).

In the supplementary floral visitation experiment, we surveyed a total of 268 floral visitors to black-eyed Susan plants from three insect taxonomic orders (Figures S2 & S3, Table S3, see Supplementary Materials for detailed results). After accounting for floral visitor taxonomic order, there was a marginally significant increase of 147% in visit duration at sodium-addition compared to control plants (χ^2^ = 3.76, df = 1, p = 0.052).

## Discussion

We show that sodium-enriched nectar can increase seed set in an animal-pollinated plant, but that this effect depended on background plant community diversity. Our findings are consistent with the ‘salty nectar hypothesis’ that plants increase the sodium concentration in their nectar to attract sodium-limited pollinators and increase seed set [6]. Additionally, we show that the effect of sodium on plant fitness is dependent on surrounding plant community diversity, with sodium-enriched plants having greater seed set in low diversity prairies but not in high diversity prairies. This context dependency highlights that while a trait such as sodium-enriched nectar may increase fitness in one habitat, it may not in another, even if the habitats are in close proximity.

To understand the potential fitness benefits of sodium-enriched nectar, we must understand sodium transport in plants. We found that black-eyed Susan (*Rudbeckia hirta*) plants had more sodium in their nectar when grown in sodium-enriched soil. Since nectaries have been gained and lost many times in angiosperms [64,65], not all nectar-producing plants may transport sodium into nectar. Our result adds another major plant family (Asteraceae) to the existing literature on sodium transport to nectar [19–21,25] (but see [30]). While sodium may have increased in both the nectar and pollen [66], the centrifuge method of nectar extraction had little to no pollen contamination, suggesting that sodium at least increased in the nectar. The ‘salty nectar hypothesis’ posits that plants transport sodium into their nectar and out of their foliar tissue to reduce herbivory while attracting pollinators [6]. For black-eyed Susan plants, foliar tissue had a lower sodium concentration than the soil, suggesting that plants were perhaps limiting sodium uptake [67,68]. The control black-eyed Susan plants had similar foliar sodium concentrations (median: 44.70 ppm, range: 27.69–118.15 ppm) to the background plant community at the site in the same year (median: 46.95 ppm, range: 22.97–59.50 ppm) (A.L. Darst, unpublished data). Notably, the site had higher background sodium concentrations than other sites in the United States Midwest, perhaps due to greater sodium deposition or differences in soil type [43,49,69].

Seed set increased in plants with sodium-enriched nectar in prairies seeded with fewer plant species. This may be because in diverse prairies, pollinators may not be able to detect or remember the locations of favored rewards in a diversity of different floral cues [33–36], although we did not directly test for this. While the increase in seed set for sodium-enriched plants in the low diversity prairies in our study is small, even a small fitness difference can be acted on by natural selection [70]. Plants that can increase the sodium in their nectar may be selected for, although this is likely limited to regions where sodium is limiting for animal pollinators. Future studies need to determine how sodium is transported into nectar, whether sodium uptake into nectar varies among individuals, and if there is a genetic component to these differences.

Seed germination was not affected by sodium additions. Seeds of other species germinated in sodium solutions have decreased germination [58,59]. However, plants may be able to limit sodium transport into seeds, or black-eyed Susan may be more sodium tolerant [37]. As an extreme example, halophyte seeds had increased germination when the mother plant was grown at a higher concentration of sodium [71]. When investigating how sodium affects plant fitness for less tolerant species, it will be important to consider both seed set and germination.

Surprisingly, we did not find an increase in floral visitation to sodium-enriched plants in our supplementary experiment, as has been reported in recent studies [24–26]. This is perhaps due to a lower sodium concentration in our natural nectar compared to studies using sodium-enriched artificial nectar [24,26] or proximity to the high diversity side of the prairie which may have affected foraging behavior [34]. Our small sample size also may have limited our ability to detect small effect sizes. However, there was a trend of longer visits to sodium-enriched flower heads. Longer visit duration may mean that floral visitors were probing each floret longer or visiting more florets, which would be expected if floral visitors preferred the sodium-enriched nectar and were foraging efficiently [7]. Longer visit durations may increase seed set through improved pollen transfer [72], but since the floral visitor surveys were conducted the year after the seed set experiment we cannot directly connect pollinator behavior to seed set in this study.

Although not the focus of this study, we observed differences in seed set and germination depending on the seeded diversity of prairies and the location of the focal plants within prairies. Control plants had significantly higher seed set in prairies seeded with more plant species. For both control and sodium-addition plants, germination rates were significantly higher in prairies seeded with more plant species and for plants closer to the middle of the prairie. This may be because pollen limitation can decrease seed set [73] and germination [74] through decreased pollen quantity and quality. Our findings are congruent with previous studies that found evidence for pollination success increasing with plant community diversity [75] and plant reproduction decreasing due to edge effects [76].

Our work demonstrates that it is not sodium alone that affects fitness; sodium interacts with the surrounding plant community to increase the fitness of sodium-enriched plants. While sodium has historically been limiting in continent interiors [11], sodium is now a common pollutant from road deicers and irrigation [43,77]. Beyond better understanding the role of sodium in plant-pollinator interactions in natural communities, our work provides insight for how novel sodium-enriched habitats could alter these interactions [25,78]. Our findings add to the growing literature of the importance of sodium, as well as other micronutrients, in ecology. Furthermore, we show that it is essential to consider context dependency in ecological questions to more fully understand the complexity of natural systems.

## Acknowledgements

Lars Brudvig conceptualized and managed the Conservation Lands Experiment (funded by NSF DEB 1547961 and 2224712). Bonnie Johnson and Cooper Pryor assisted with field work. Jamie Smith grew black-eyed Susans for floral visitation surveys. Lindsey Kemmerling, Mark Hammond, Holly Vander Stel, Jeff Conner, Kate Shaw, and Robin Waterman provided advice for growing plants, extracting nectar, and measuring seed set. Nate Sanders and Paul CaraDonna provided helpful feedback. Sodium analysis was performed by Keith MacRenaris at the Michigan State University Quantitative Bio Element Analysis and Mapping Center, which is supported by the Office of the Vice President for Research and Innovation, the Colleges of Natural Sciences, Human Medicine, Osteopathic Medicine, Veterinary Medicine, Engineering, Agriculture and Natural Resources with additional funding provided by the National Research Resource for Quantitative Mapping in the Life Sciences by the National Institute of General Medical Sciences under grant #P41GM135018. This is W.K. Kellogg Biological Station contribution no. ####.

## Author contributions

A.L.D.: conceptualization, data curation, formal analysis, funding acquisition, investigation, methodology, visualization, writing—original draft, writing—review and editing; M.E.L.: conceptualization, investigation, methodology, writing—review and editing; N.M.H.: conceptualization, supervision, writing—review and editing.

## Data accessibility statement

Data and code will be archived at Zenodo upon acceptance.

