## Supplementary Material for "Sodium-enriched nectar increases seed set in low diversity prairies"

**Supplementary Methods**

*Floral visitation surveys*

In a supplemental experiment to further understand how sodium-enriched nectar affected plant fitness, we conducted floral visitation surveys the following summer to explore visitor preference (see Supplementary Methods for additional methods and results). Plants used for floral visitation surveys were grown from the same seed stock as those used in germination and seed set trials and were treated using the same methods. Between June 2, 2025 and June 16, 2025, we surveyed four pairs of sodium-addition and control plants daily for ten days to investigate floral visitor preference. Each plant pair was surveyed between 7–10 times. Surveys only occurred on sunny days above 15.5°C without rain. We placed plant pairs in the center of one CLE prairie, between the high and low diversity sides of the field. For twenty minutes, two surveyors, one per plant, counted and identified floral visitors, timed visit duration, counted the number of flower heads visited, and noted whether floral visitors moved between the two plants. Coleopterans (n = 3) and hemipterans (n = 2) visited the flowers but were not included in analyses, as they were unlikely to be acting as pollinators [1,2]. Additionally, surveys in which surveyors were unable to time all the floral visitors were not included in analyses of visit duration. The surveyors alternated between surveying the sodium-addition or control plant, the plant location during the survey, and the order in which plant pairs were surveyed.

To investigate how sodium affected floral visitation, we first fit three generalized linear mixed effects models in R package 'glmmTMB' v.1.1.14 [3]. We fit floral visitor abundance, number of flower heads visited, or visit duration as a function of sodium treatment and the number of flower heads on the plant, with plant nested in pair as a random intercept to account for repeated sampling. Because different floral visitor groups may not all respond to sodium in the same way [4], we repeated analyses with an interaction between sodium treatment and floral visitor taxonomic order. The best fitting and most parsimonious models (dAICc > 2, Table S1) used a negative binomial distribution for floral visitor abundance and number of flower heads visited, and a zero inflated negative binomial distribution for visit duration.

**Supplementary Results**

*Floral visitation surveys*

We surveyed a total of 268 floral visitors to black-eyed Susan plants from three insect taxonomic orders (Table S3). The majority of floral visitors were in the family Halictidae ($n \geq 195$). Sodium-addition did not affect total floral visitor abundance ($\chi^2 = 0.03$, df = 1, p = 0.87), total flower heads visited ($\chi^2 = 0.02$, df = 1, p = 0.88), or duration visited ($\chi^2 = 0.08$, df = 1, p = 0.78, Fig. S1, Table S4). Separating floral visitors into Diptera, Hymenoptera, and Lepidoptera revealed subtle differences in visitation by sodium treatment (Fig. 2, Table S5, Table S6). Sodium-addition did not affect floral visitor abundance ($\chi^2 = 0.13$, df = 1, p = 0.71) or number of flower heads visited ($\chi^2 = 0.02$, df = 1, p = 0.90), and this effect did not differ by taxonomic order for floral visitor abundance ($\chi^2 = 0.59$, df = 2, p = 0.75) or number of flower heads visited ($\chi^2 = 2.79$, df = 2, p = 0.25). However, after accounting for floral visitor taxonomic order, there was a marginally significant increase of 147% in visit duration at sodium-addition compared to control plants ($\chi^2 = 3.76$, df = 1, p = 0.052). The interaction between sodium treatment and floral visitor taxonomic order was not significant ($\chi^2 = 4.03$, df = 2, p = 0.13). Additionally, of the seven individual floral visitors observed foraging on both plants in a pair, six bees switched from foraging on the control plant to the sodium-addition plant.

**Supplementary Figures**

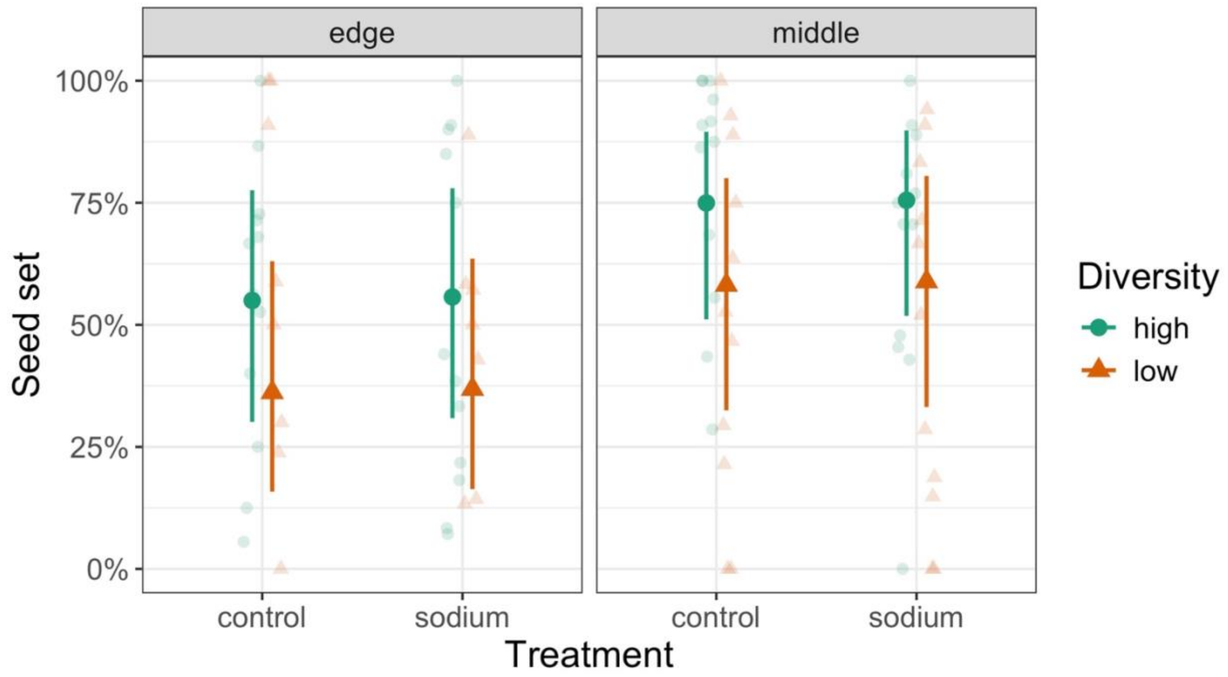


Figure S1. Germination rates for sodium-addition and control black-eyed Susan plants in prairies with high and low seeded diversity and in exclosures near the edge and middle of prairies. Solid points represent the estimated marginal means. Error bars are 95% confidence intervals. Translucent points are raw, jittered data.

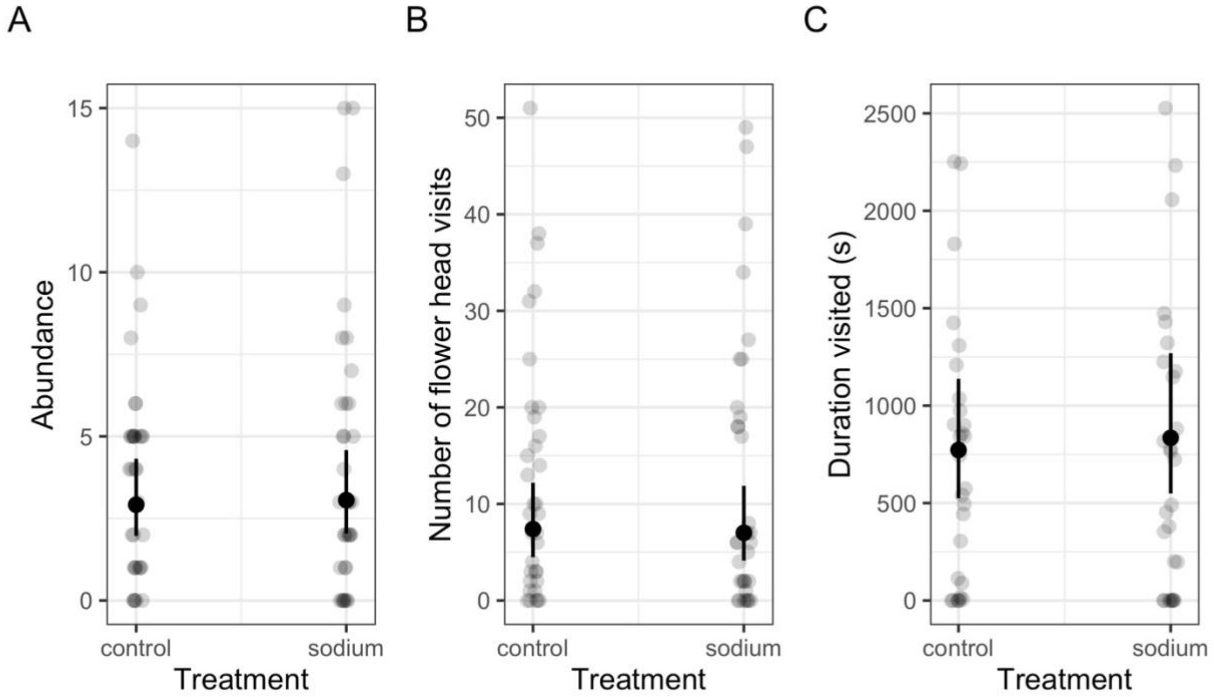


Figure S2. Floral visitor A) abundance, B) number of flowers visited, and C) duration visited for sodium-addition and control black-eyed Susan plants. Solid points represent the estimated marginal means. Error bars are 95% confidence intervals. Translucent points are raw, jittered data.

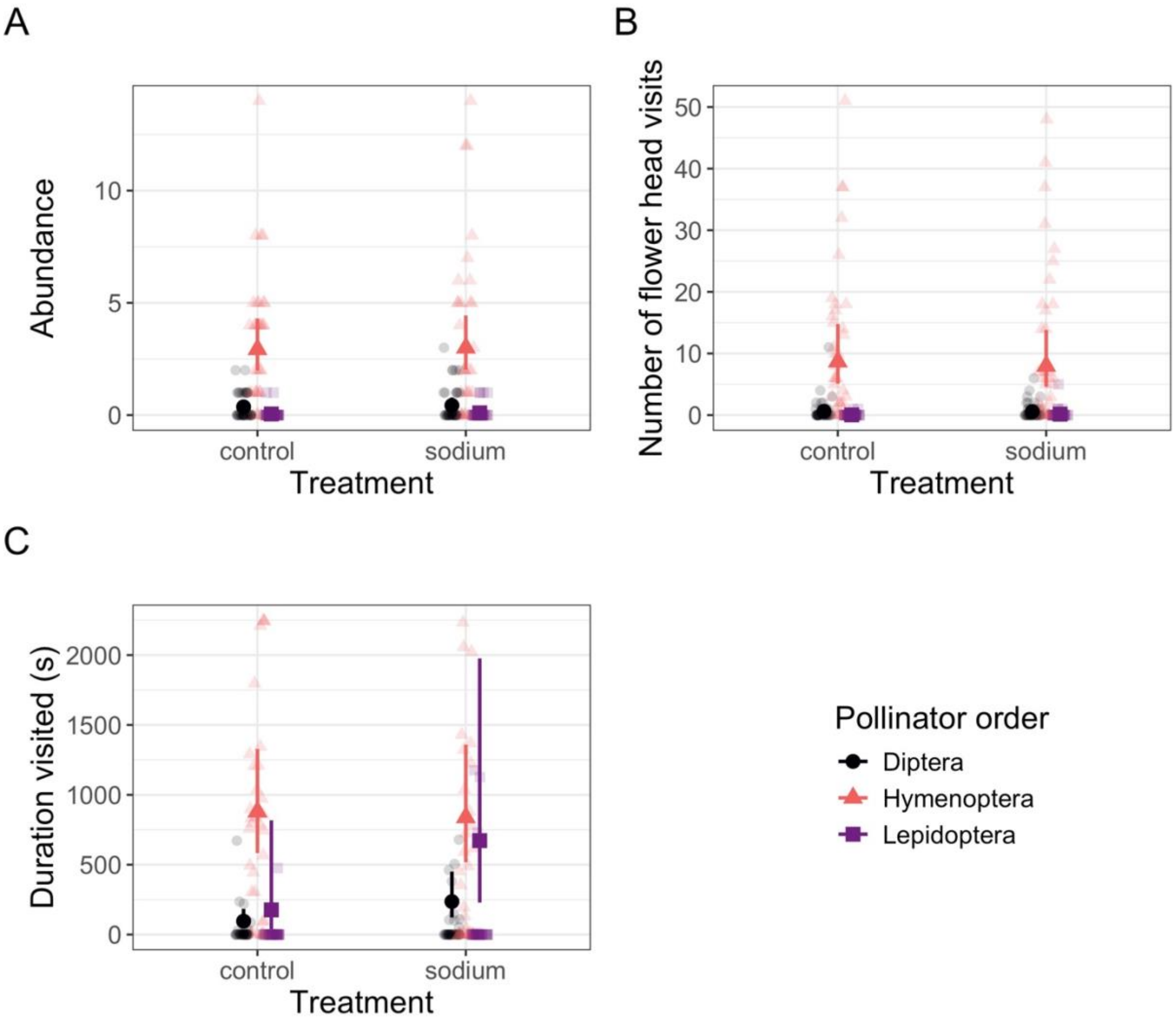


Figure S3. Floral visitor taxonomic order A) abundance, B) number of flower heads visited, and C) duration visited for sodium-addition and control black-eyed Susan plants. Solid points represent the estimated marginal means. Error bars are 95% confidence intervals. Translucent points are raw, jittered data.

**Supplementary Tables**

Table S1. Model comparison of distributions for floral visitor abundance, number of flower heads visited, or visit duration using dAICc. Models without treatment (sodium/control) were excluded from comparisons. The table is sorted by dAICc. The bolded model had the lowest dAICc. NC = non-convergence

| Response | Distribution | Zero-inflation | dAICc | df |
|---|---|---|---|---|
| **Abundance** | **Negative binomial: quadratic** | **no** | **N/A** | **10** |
| | Negative binomial: quadratic | yes | NC | NC |
| | Poisson | no | NC | NC |
| | Poisson | yes | NC | NC |
| **Number of flower head visits** | **Negative binomial: quadratic** | **no** | **0.0** | **10** |
| | Negative binomial: quadratic | yes | 2.2 | 11 |
| | Poisson | yes | 251.0 | 10 |
| | Poisson | no | NC | NC |
| **Duration visited (s)** | **Negative binomial: quadratic** | **yes** | **0.0** | **11** |
| | Negative binomial: quadratic | no | 43.1 | 10 |
| | Poisson | yes | 24935.2 | 10 |
| | Poisson | no | NC | NC |

Table S2. Model comparison for seed set and germination using dAICc and log-likelihood. Models without treatment (sodium/control) were excluded from comparisons. The table is sorted by dAICc. The bolded model had the lowest dAICc. NC = non-convergence

| Response | Model terms | dAICc | log-likelihood | df |
|---|---|---|---|---|
| **Seed set** | **Treatment*Diversity** | **0.0** | **-337.0** | **6** |
| | Treatment*Diversity + Location | 1.4 | -336.6 | 7 |
| | Treatment + Diversity | 8.2 | -342.3 | 5 |
| | Treatment + Diversity*Location | 8.3 | -340.0 | 7 |
| | Treatment | 9.3 | -343.9 | 4 |
| | Treatment + Diversity + Location | 9.6 | -341.8 | 6 |
| | Treatment + Location | 10.8 | -343.6 | 5 |
| | Treatment*Diversity *Location | NC | NC | NC |
| **Germination rate** | **Treatment + Diversity + Location** | **0.0** | **-224.6** | **8** |
| | Treatment + Diversity*Location | 0.9 | -223.8 | 9 |
| | Treatment*Diversity + Location | 2.4 | -224.6 | 9 |
| | Treatment + Diversity | 2.6 | -227.1 | 7 |
| | Treatment | 3.0 | -228.5 | 6 |
| | Treatment*Diversity | 5.0 | -227.1 | 8 |
| | Treatment + Location | NC | NC | NC |
| | Treatment*Diversity *Location | NC | NC | NC |

Table S3. Number of floral visitors to sodium-addition and control black-eyed Susans (*Rudbeckia hirta*). Bees were identified to morphogroup [5].

| Floral visitor order | Floral visitor lowest identity | Floral visitor family | N control | N sodium | N total |
|---|---|---|---|---|---|
| Hymenoptera | Striped sweat bee | Halictidae | 59 | 68 | 127 |
| | Green sweat bee | Halictidae | 31 | 37 | 68 |
| | Tiny dark bee | Halictidae, Apidae, or Colletidae | 10 | 9 | 19 |
| | Medium dark bee | Andrenidae | 8 | 3 | 11 |
| | Honey bee (*Apis mellifera*) | Apidae | 4 | 1 | 5 |
| | Striped hairy belly bee | Megachilidae | 1 | 0 | 1 |
| | | **TOTAL** | **113** | **118** | **231** |
| Diptera | Hoverfly | Syrphidae | 7 | 9 | 16 |
| | Dronefly (Eristalini) | Syrphidae | 4 | 5 | 9 |
| | Unknown fly | | 3 | 3 | 6 |
| | | **TOTAL** | **14** | **17** | **31** |
| Lepidoptera | Plume moth | Pterophoridae | 1 | 2 | 3 |
| | Yellow collared scape moth (*Cisseps fulvicollis*) | Erebidae | 1 | 2 | 3 |
| | | **TOTAL** | **2** | **4** | **6** |

Table S4. Model estimates and type II ANOVA statistics for floral visitor abundance, number of flower heads visited, or visit duration as a function of the number of flowers on the plant and treatment (sodium/control) with plant nested in pair as a random intercept.

| Response | Predictors | β | SE | $\chi^2_{df = 1}$ | p-value |
|---|---|---|---|---|---|
| Abundance | # flowers | 0.076 | 0.016 | 22.080 | < 0.0001 |
| | Treatment (sodium) | 0.047 | 0.280 | 0.028 | 0.866 |
| Number of flower head visits | # flowers | 0.127 | 0.023 | 31.175 | < 0.0001 |
| | Treatment (sodium) | -0.054 | 0.365 | 0.022 | 0.882 |
| Duration visited (s) | # flowers | 0.037 | 0.023 | 2.544 | 0.111 |
| | Treatment (sodium) | 0.077 | 0.274 | 0.080 | 0.777 |

Table S5. Model estimates for floral visitor abundance, number of flower heads visited, or visit duration as a function of the number of flowers on the plant and the interaction between treatment (sodium/control) and floral visitor order with plant nested in pair as a random intercept.

| Response | Predictors | β | SE |
|---|---|---|---|
| Abundance | Intercept | -1.841 | 0.379 |
| | # flowers | 0.071 | 0.014 |
| | Treatment (sodium) | 0.161 | 0.443 |
| | Order (Hymenoptera) | 2.076 | 0.330 |
| | Order (Lepidoptera) | -1.991 | 0.776 |
| | Treatment (sodium):Order (Hymenoptera) | -0.137 | 0.450 |
| | Treatment (sodium):Order (Lepidoptera) | 0.527 | 0.969 |
| Number of flower head visits | Intercept | -1.860 | 0.439 |
| | # flowers | 0.116 | 0.018 |
| | Treatment (sodium) | -0.064 | 0.502 |
| | Order (Hymenoptera) | 2.648 | 0.361 |
| | Order (Lepidoptera) | -2.786 | 0.818 |
| | Treatment (sodium):Order (Hymenoptera) | -0.021 | 0.505 |
| | Treatment (sodium):Order (Lepidoptera) | 1.493 | 0.963 |
| Duration visited (s) | Intercept | 4.233 | 0.443 |
| | # flowers | 0.029 | 0.018 |
| | Treatment (sodium) | 0.902 | 0.465 |
| | Order (Hymenoptera) | 2.218 | 0.383 |
| | Order (Lepidoptera) | 0.614 | 0.832 |
| | Treatment (sodium):Order (Hymenoptera) | -0.951 | 0.563 |
| | Treatment (sodium):Order (Lepidoptera) | 0.433 | 1.047 |

Table S6. Type III ANOVA statistics for floral visitor abundance, number of flower heads visited, or visit duration as a function of the number of flowers on the plant and the interaction between treatment (sodium/control) and floral visitor order with plant nested in pair as a random intercept.

| Response | Predictors | df | $\chi^2$ | p-value |
|---|---|---|---|---|
| Abundance | # flowers | 1 | 26.914 | < 0.0001 |
| | Treatment | 1 | 0.131 | 0.717 |
| | Order | 2 | 64.381 | < 0.0001 |
| | Treatment:Order | 2 | 0.587 | 0.746 |
| Number of flower head visits | # flowers | 1 | 39.268 | < 0.0001 |
| | Treatment | 1 | 0.016 | 0.899 |
| | Order | 2 | 86.446 | < 0.0001 |
| | Treatment:Order | 2 | 2.792 | 0.248 |
| Duration visited (s) | # flowers | 1 | 2.559 | 0.110 |
| | Treatment | 1 | 3.764 | 0.052 |
| | Order | 2 | 34.645 | < 0.0001 |
| | Treatment:Order | 2 | 4.026 | 0.134 |

Table S7. Type III ANOVA statistics for seed set and germination rate. We fit seed set as a function of the interaction between seeded diversity and treatment with pair nested in CLE site as a random intercept. We fit germination rate as a function of treatment (sodium/control), seeded diversity, and exclosure location with the random intercepts pair nested in CLE site and growth chamber shelf nested in growth chamber.

| Response | Predictors | $\chi^2_{df=1}$ | p-value |
|---|---|---|---|
| Seed set | Treatment | 3.593 | 0.058 |
| | Diversity | 7.336 | 0.007 |
| | Treatment:Diversity | 10.506 | 0.001 |
| Germination rate | Treatment | 0.034 | 0.854 |
| | Diversity | 3.953 | 0.047 |
| | Location | 5.303 | 0.021 |

Table S8. Pairwise comparisons of estimated marginal means for seed set. Significant comparisons are bolded (α < 0.05).

| Contrast | Odds ratio | SE | z-ratio | p-value |
|---|---|---|---|---|
| **high control / low control** | **2.184** | **0.630** | **2.708** | **0.034** |
| high control / high sodium | 1.233 | 0.136 | 1.895 | 0.230 |
| high control / low sodium | 1.564 | 0.450 | 1.555 | 0.405 |
| high sodium / low sodium | 1.268 | 0.365 | 0.826 | 0.842 |
| **low sodium / low control** | **1.397** | **0.176** | **2.649** | **0.040** |
| high sodium / low control | 1.771 | 0.511 | 1.982 | 0.195 |
